# Base-pair resolution detection of transcription factor binding site by deep deconvolutional network

**DOI:** 10.1101/254508

**Authors:** Sirajul Salekin, Jianqiu (Michelle) Zhang, Yufei Huang

**Affiliations:** Electrical and Computer Engineering Department, University of Texas at San Antonio, 1 UTSA Circle, San Antonio, TX 78249, USA.; Department of XXXXXXX, Address XXXX etc.

## Abstract

**Motivation:** Transcription factor (TF) binds to the promoter region of a gene to control gene expression. Identifying precise transcription factor binding sites (TFBS) is essential for understanding the detailed mechanisms of TF mediated gene regulation. However, there is a shortage of computational approach that can deliver single base pair (bp) resolution prediction of TFBS.

**Results:** In this paper, we propose DeepSNR, a Deep Learning algorithm for predicting transcription factor binding location at Single Nucleotide Resolution de novo from DNA sequence. DeepSNR adopts a novel deconvolutional network (deconvNet) model and is inspired by the similarity to image segmentation by deconvNet. The proposed deconvNet architecture is constructed on top of ‘Deep-Bind’ and we trained the entire model using TF specific data from ChIP-exonuclease (ChIP-exo) experiments. DeepSNR has been shown to outperform motif search based methods for several evaluation metrics. We have also demonstrated the usefulness of DeepSNR in the regulatory analysis of TFBS as well as in improving the TFBS prediction specificity using ChIP-seq data.

**Availability:** DeepSNR is available open source in the GitHub repository (https://github.com/sirajulsalekin/DeepSNR)

## 1 Introduction

Transcription factor binding sites are specific DNA sequences that control gene expression through interaction with transcription factor proteins. Revealing the dynamic regulatory systems by transcription factors (TFs) signifies one of the major challenges in biological research. Precise mapping of TFBSs on a genomic scale plays a pivotal role in delineating transcription regulatory network and remains a long sought goal in genomic annotations (J.-t. Guo, Lofgren, & Farrel, 2014; Salekin, Bari, Raphael, Forsthuber, & Zhang, 2016, 2017). Chromatin immunoprecipitation (ChIP) that yields a set of statistically enriched high occupancy binding regions is the most widely used method to recognize protein-DNA binding locations (Peng, Alekseyenko, Larschan, Kuroda, & Park, 2007; Tuteja, White, Schug, & Kaestner, 2009). However, incongruent size of randomly clipped DNA fragments in ChIP technology largely limits the resolution of ChIP-seq data. To overcome this limit, ChIP-exo technique was developed that uses λ phage exonuclease to digest the 5′ end of TF-unbound DNA after ChIP (Rhee & Pugh, 2011). In ChIP-exo, λ exonuclease digestion leaves homogenous 5′ ends of DNA fragments at the actual two boundaries of TFBS, and after sequencing and mapping reads to the reference genome two borders of TFBS could be defined. The λ exonuclease treatment augments signal-to-noise ratio by eliminating unwanted DNA, which allows the discovery of low affinity binding sites.

With the advent of rapidly increasing genomic sequences, sequence-based computational methods have been developed and proven to be valuable in predicting TFBS (J.-t. Guo et al., 2014; Stormo, 2000). The computational methods generally scrutinizes user provided input sequences in order to identify TF binding motifs that are statistically overrepresented in binding sites with respect to background sequence. Predicting the binding location based on motif suffers from several shortcomings. First, motifs are typically short 10-15 bp sequences and therefore prediction using binding motifs is unlikely to generate predictions with high specificity. Moreover, motifs represent only the enriched binding sequence patterns and thus cannot explain all possible bindings of a TF. Finally, even if the motif search methods succeed in determining the anchor position of a putative binding site, they cannot predict the actual width of TFBS. The specificity of protein-DNA binding does not depend only on DNA sequence, but it also depends on the 3D structures of DNA and TF protein macromolecules (Rohs et al., 2010) which explains the failure of motif searches in predicting true TFBS.

To enable precise prediction of TFBS, we designed in this paper a deep learning based model called DeepSNR. Deep learning, the most active field in machine learning, has been proven to achieve record-breaking performances in image and speech recognition (Graves, Mohamed, & Hinton, 2013; Zeiler & Fergus, 2014), natural language understanding (Sutskever, Vinyals, & Le, 2014; Xiong, Merity, & Socher, 2016), and most recently, in computational biology (Alipanahi, Delong, Weirauch, & Frey, 2015; Hassanzadeh & Wang, 2016; Quang & Xie, 2016; Zhou & Troyanskaya, 2015). The two recent methods, DeepBind (Alipanahi et al., 2015) and DeepSEA (Zhou & Troyanskaya, 2015), successfully applied deep learning to model the sequence specificity of TF binding with a performance superior to the best existing shallow learning methods. Convolutional neural network (CNN) was adopted by these methods to capture the features essential for accurate characterization of motifs for target TFs. DeeperBind (Hassanzadeh & Wang, 2016) and DanQ (Quang & Xie, 2016) employed recurrent neural network (RNN) along with CNN to learn the spatial dependencies of detected motifs and yielded improved prediction performance in comparison to DeepBind and DeepSEA respectively. In spite of their success in determining the presence of binding site in a given DNA sequence, these approaches cannot report the precise binding location. Our proposed method intends to bridge the gap by identifying transcription factor binding location at single nucleotide resolution from 100 bp long input DNA sequence that is known to contain TFBS (e.g. ChIP-seq regions). DeepSNR is inspired by the similarity between ascertaining the TF binding location from 100 bp long sequences and image segmentation method. Similar to pixel-level image segmentation where each pixel is categorized as belonging to target object (e.g. dog, car, human) or background, DeepSNR classifies each nucleotide in a DNA sequence as putative binding site or background sequence and thereby achieves base pair resolution prediction. Recently, deconvNet (Noh, Hong, & Han, 2015) has achieved remarkable success in semantic image segmentation that aims to predict a category label for every image pixel. In that study, the authors built the deconvNet on top of the CNN obtained from VGG 16-layer net. Comparatively, the multi-layer deconvolution network in DeepSNR is composed of convolution layers adopted from DeepBind, deconvolution, unpooling, and rectified linear unit (ReLU) layers (Fig. 1). Instead of relying on the similarity of binding sequences for deriving the binding preference of a transcription factor, DeepSNR accurately captures the inherent complex interactions between TF and DNA and thus enables it to precisely locate the binding site.

The entire deconvNet is trained using the data generated by ChIP-exo experiment and can be applied to individual sequences to pinpoint the TFBS location. When tested, DeepSNR attained outstanding result that substantially surpasses binding motif based algorithms in terms of precision, recall, F-Score and IoU. For instance, the trained DeepSNR model for CTCF achieved 83% median F-Score over 19,600 test sequences while MatInspector managed to record only 58%. We further discovered that the trained model automatically detects the location of motif sequence in pursuit of identifying binding site. When we applied DeepSNR onChIP-seq data, it rendered us with unique display of distribution of TF binding motif over the ChIP-seq binding area which has been possible to visualize because of the base-pair resolution prediction of DeepSNR (Supplementary figure S2). We have also demonstrated the capacity of DeepSNR in improving the specificity of ChIP-seq peak calling results by an independent motif enrichment analysis that confirms the presence of highly enriched motif sequence in DeepSNR predicted binding region (Table 1). Moreover, the capability of DeepSNR in pinpointing the motif sequence in ChIP-seq data makes the model suitable for playing a role in regulatory analysis of TFBS.

**Fig. 1.**
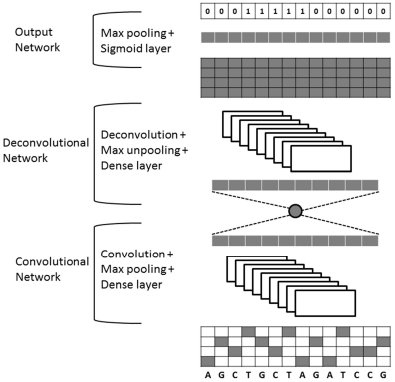
DeepSNR model architecture

## 2 Methods

This section discusses the architecture of DeepSNR model and describes the overall TFBS discovery algorithm.

### 2.1 Model Design/Architecture

Fig. 1 illustrates the detailed configuration of DeepSNR which is composed of three parts – convolutional, deconvolutional and output networks. The input of DeepSNR is a one-hot encoded 100 bp long DNA sequence that is known to contain a binding site of the TF of interest. While the convolutional network corresponds to feature extractor that learns the inherent features imperative for TF-DNA binding, the deconvolution network is a shape generator that locates the binding site using the feature extracted from the convolution network. The output network of the model is used to generate a 100 bit binary sequence, indicating whether each nucleotide belongs to binding site (1) or background sequence (0).

The convolutional part of DeepSNR model is a replica of DeepBind (Alipanahi et al., 2015) network, which consists of one convolutional layer, followed by rectification and pooling operation, and one fully connected network (FCN) augmented at the end to transform feature vectors into a scalar binding score. Our deconvolution network is a mirrored version of the convolution network, and has a series of unpooling, deconvolution, and rectification layers. Contrary to convolutional network that reduces the size of activations through feedforward step; deconvolutional network enlarges the activations through combinations of unpooling and deconvolution operations. The last layer, the sigmoid output layer, makes binary predictions for each of the 100 nucleotides. We get the maximum for each base pair over the output of deconvolution layer before employing the sigmoid function and then we apply a threshold to map the final output to 0/1.

To implement max-unpooling and deconvolution operations, we followed the similar procedure described in (Noh et al., 2015; Zeiler & Fergus, 2014). The model records the position of maximum activation while performing the pooling operation and later this information is used in unpooling procedure to assign each stimulus back into appropriate location. The unpooling layer is especially important because retaining the place of maxima assists in capturing the binding motif and the binding site associated 3D contextual information and proves to be critical for precise localization of TFBS. The output of deconvolution layer associates a single input activation with multiple outputs, as illustrated in (Noh et al., 2015). The deconvolution layer employed in our model is fundamentally the reverse operation of convolution and used to learn the shape details of TFBS. Integrating this layer in the architecture helps DeepSNR to capture the overall breadth of a binding site, thus improving the completeness of the model.

### 2.2 Training the DeepSNR Model

The entire deconvolutional network is comprised of seven layers and contains a lot of associated parameters. In addition, the parameter search space for predicting binding location is enormous because TF-DNA binding is a very complicated phenomenon depending on DNA sequence, 3D structure of DNA and TF protein and their intrinsic complex interactions. Therefore, we trained DeepSNR in two stages as in (Noh et al., 2015), so that the model progressively learns the essential features to recognize TFBS and tunes to optimum set of parameters. For the first stage of training, we constructed the training set such that the binding sites were placed at the center of 100 bp long input sequence. By doing so, we limited the search space significantly and forced the model to learn the intricate details of TF-DNA binding. We initialized the weights in convolutional network using DeepBind pre-trained for specific transcription factor, while the weights in deconvolutional network were initialized with random samples from zero-mean Gaussians. Initializing the weights with DeepBind convolutional network is very important because it assists the model to converge with minimum iterations and mitigates vanishing gradient problem. In the second stage, we imposed the model with more challenging training samples by placing the binding site in random locations within input sequence as described in the next section. Weights learned from the first stage of training were used to initialize all the layers in this stage and they were fine-tuned making the network robust to TF binding location. Another major challenge in training a deep network is the modification of weight distributions due to the parameter updates of preceding layers which amplifies through propagation across layers (Ioffe & Szegedy, 2015). Hence, we performed batch normalization at the output of convolutional and deconvolutional layer to better optimize our network.

To train the model, we minimized the sigmoid cross-entropy loss which essentially leads to binary logistic regression. The standard stochastic gradient descent was employed for optimization, where the learning rate was set to 0.01. The stochastic gradient descent (SGD) method estimates the training objective gradient using only a subset of training examples. The batch size determines how many training pairs to sample for each parameter update step. In our implementation, the batch size was equal to 100 samples. The network converges after approximately 15K and 20K iterations respectively in first and second stage and the training takes less than an hour in a single computer with 12G memory. We implemented the proposed network based on tensorflow. Lastly, a threshold was set at the output layer of DeepSNR architecture to return binary outcome that indicates whether a nucleotide belongs to binding site or not. We learned the threshold using validation set such that average F-Score (supplementary section S1) is maximized over the whole set and then, applied it on the test set for performance evaluation of DeepSNR.

### 2.3 Data for Training DeepSNR

We employed published human CTCF ChIP-exo data (accession number: SRA044886) for training and testing the proposed DeepSNR. Engaging the highly sensitive ChIP-exo experimental data is imperative to train DeepSNR because it aids our model to learn essential contextual information to precisely locate TFBS. We applied MACE (Wang et al., 2014) to the ChIP-exo data to identify genome-wide mapping of CTCF binding sites. MACE identified total 110,183 CTCF binding sites across the whole genome. After investigating the size distribution of those sites, we observed that 59,425 sites’ width was equal to 49 bp in accordance with the previously studied results (Rhee & Pugh, 2011; Wang et al., 2014). Hence, we utilized these 49 bp long TFBSs and added 51 bp flanking regions from two sides to make each sample 100 bp long. The flanking regions provide extra contextual information about TFBS to the model. The test set contains 19,600 randomly selected samples and the rest of the samples were used for training (34,925 sites) and validation (5000 sites). The training, testing and validation samples are strictly non-overlapping.

Each training sample consists of a 100 bp sequence from the human hg19 reference genome and is paired with a label vector of same size indicating TFBS location. To construct training samples, we used the same training and validation set in both training stages with only one difference. While in the first stage, we placed the 49 bp long binding region at position 26-74 bp of a 100 bp input sequence for all the training and validation samples, the binding sites were positioned contiguously anywhere between 10-90 bp for the second stage of training. For the 19,600 samples of test set, the input sequences were generated in the same procedure as second stage and we utilized it to assess the performance of DeepSNR only after both stages of training were completed.

We also trained separate DeepSNR models to predict binding location of androgen receptor (AR) and glucocorticoid receptor (GR) transcription factors. Table S1 summarizes the dataset information of these transcription factors.

## 3 Results

MatInspector (Cartharius et al., 2005) and MATCH (Kel et al., 2003) are two of the widely used DNA sequence based computational approaches for determining the location of TFBS. These methods scan input DNA sequences using position weight matrix (PWM) model of the desired transcription factor and assign matrix similarity score (MSS) for each K-mer. After assigning MSS, a cut-off threshold is set to decide putative binding site. In order to evaluate the effectiveness and efficiency of our proposed approach, we compare the performance of DeepSNR with MatInspector on the CTCF dataset derived from ChIP-exo experiment. MATCH was not considered for the comparison analysis because MatInspector and MATCH perform quite similarly because both the methods rely on motif search for predicting TFBS. There are several methods that also provide base pair resolution prediction of TFBS using ChIP-seq data such as GEM (Y. Guo, Mahony, & Gifford, 2012), PeakZilla (Bardet et al., 2013) and PICS (Zhang et al., 2011). However, all of these methods require ChIP-seq read distribution information to predict binding location. Since, DeepSNR is not a peak calling method and is designed to identify TF binding region using DNA sequences only; we did not compare the prediction performance with those methods.

### 3.1 Performance Evaluation Scheme

The goal of the proposed algorithm is to precisely identify the TFBS location from 100 bp long DNA sequence at single bp resolution. That is, for each nucleotide of input sequence, we aim to determine whether the base-pair categorizes to putative binding site or contextual sequence. Hence, we employed Intersection-over-Union (IoU) between ground-truth and predicted location as one of the evaluation metric to assess performance. IoU is very popular in the field of pixel-level image segmentation since it discerns a proposed solution with respect to the ground truth in perceptually meaningful way. We have also compared the performance of DeepSNR and MatInspector in terms of precision, recall and F-Score (supplementary section S1).

To assess the efficacy of DeepSNR in regard to the metrics mentioned above, we distinctly tested the performance of each independently trained DeepSNR model for CTCF, GR and AR transcription factors. Since, our method yields 1/0 for each nucleotide in a sequence, we calculated precision, recall, F-Score and IoU for each of the input sequences individually for further analysis. For CTCF, the whole test set which comprises 19,600 distinct DNA sequences of 100 bp length each was used to estimate the performance of DeepSNR. On the other hand, the performance of MatInspector was assessed slightly differently. When we run MatInspector over 19,600 test sequences with all the parameters set to optimized values as determined by the algorithm, we found that the method was able to detect TFBS only in 2,942 sequences. We investigated the missed sequences using MEME (Machanick & Bailey, 2011) and FIMO (Grant, Bailey, & Noble, 2011) and learned that those sequences contain highly enriched CTCF motifs, albeit a degenerate version. This maybe explains the failure of MatInspector in discovering any binding site for those sequences. However, we used these 2,942 sequences to measure the performance of MatInspector and compared the results with DeepSNR which were calculated based on 19,600 test sequences, though it is advantageous for MatInspector algorithm.

### 3.2 Performance Analysis of DeepSNR and MatInspector

In this section, we comprehensively analyze the performance of DeepSNR and MatInspector using the ground truth TFBSs locations derived from CTCF ChIP-exo dataset. Since, ChIP-exo reports TFBS location at single nucleotide sensitivity, using it as ground truth helps eliminating any ambiguity in performance comparison between different methods. The box plots in Fig. 2(a) show median values of evaluation metrics when calculated over all sequences as described in the previous section. It is clear that DeepSNR outperforms MatInspector to a large extent. The median recall of DeepSNR over all test sequences is 91% and it achieves sensitivity greater than 98% for at least 25% of the sequences under consideration. On the other hand, the best recall recorded by MatInspector for any sequence is merely 55%. The large difference in recall score between two methods emphasizes that DeepSNR is very sensitive in locating TFBS at base-pair resolution and it can successfully predict the total width of binding site instead of identifying just the anchor position.

**Fig. 2.**
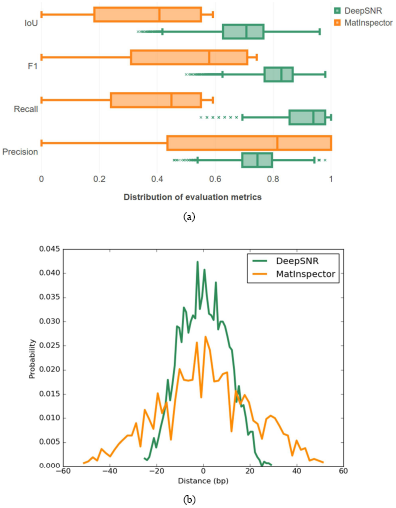
(a) Performance comparison of DeepSNR and MatInspector, (b) Distribution of the distance of center nucleotide for DeepSNR and MatInspector from that of ground truth binding sites

The median specificity of our system against false positives seems to be lower than MatInspector. However, DeepSNR demonstrates a precision higher than 73% for at least 10,000 test sequences whereas the median precision of MatInspector is 81% despite the fact that the statistics was measured across 2,947 sequences only. Combining the precision and recall evaluation metrics in the F-Score measure shows that DeepSNR significantly improved the prediction performance of locating TFBSs by 25% compared with MatInspector. Furthermore, DeepSNR accomplishes the F-Score as high as 98% for some of the test sequences. Finally, the DeepSNR has IoU greater than 68% on half of the test sequences while MatInspector achieves a median IoU score of only 40%. The higher IoU of DeepSNR indicates that the method is sensitive enough to recognize nucleotides belonging to true TFBSs without conceding to precision, which is a remarkable feat.

This pattern extends to the remaining transcription factors as DeepSNR outperforms MatInpector by 23.8% and 48.5% respective improvement of IoU for GR and AR (Supplementary Fig S1). The center position of TFBS is also important for downstream analysis. Hence, we also investigated distance between the centers for predicted sites and the centers of ground truth binding sites of test dataset. As evident from Fig. 2(b), the distance density plot for DeepSNR predicted center binding region is highly focused at the vicinity of zero in comparison to MatInspector. We found that DeepSNR displayed distance mean 0.69 bp and SD 10.23 bp while the mean and SD for MatInpector were found to be 1.95 bp and 20.37 bp respectively (p-value = 2e-5).The narrower peak of DeepSNR illustrates that the center nucleotide of predicted binding sites mostly coincide with the center of true binding site. Overall, these results demonstrate that the proposed deep learning model successfully captures TF-DNA binding interactions and improves the prediction of binding location.

### 3.3 DeepSNR precisely locates binding motif

To rigorously understand the significance of the results predicted by the trained DeepSNR model, we investigated the output of max-pooling layer at output network for 19,600 CTCF test sequences. The max-pooling layer yields 100 scaler numbers (scores) upon which the sigmoid function and a threshold is applied to produce binary outcome corresponding to each base-pair of an input sequence. We surmised that the base-pairs having higher scores may indicate biologically significant nucleotides for transcription factor (TF) binding. Hence, mutation map scheme (Alipanahi et al., 2015) was deployed to identify the most significant nucleotide (Ntms) for TF binding (see supplementary section S2). Then we assessed distance of Ntms from the nucleotide having maximum score (Ntmax) as computed by DeepSNR or (Ntms – Ntmax). Figure 3a shows the histogram of distances measured across 19,600 CTCF test sequences. It is evident from the histogram that in a large portion (57%) of test sequences the nucleotide whose mutation predominantly impacts the binding affinity is positioned at 10-18 bp apart from Ntmax. Interestingly, it has been shown in a previous study that nucleotides 4-8 and 10-18 within core motif of a CTCF binding site (CBS) are the most critical determinant for CTCF binding (Plasschaert et al., 2014; Renda et al., 2007). Since the nucleotides maximally influencing the CTCF binding because of point mutations are commonly placed at a distance of 10-18 bp from Ntmax, we deduced that the base achieving maximum score according to DeepSNR (Ntmax) marks the first nucleotide of CTCF binding motif in CBS.

**Fig. 3.**
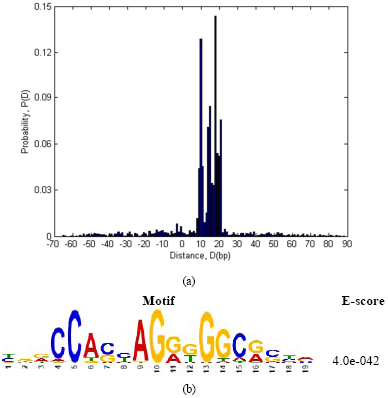
(a) Probability distribution of distance of the most significant nucleotide (Ntms) for TF binding identified by DeepBind from the nucleotide having maximum score (Ntmax) according to DeepSNR. (b) CTCF motif found as highly present after performing motif enrichment analysis on all test sequences for very short region, only 20 bp downstream from Ntmax

To further validate our implication that Ntmax truly indicates the first nucleotide of CTCF core motif within CBS, we performed motif enrichment analysis using MEME within 20 bp downstream from Ntmax. CTCF binding motifs are known to be ∼20 bp long (Plasschaert et al., 2014; Rhee & Pugh, 2011). Hence, we selected 20 bp only to impose the most stringent criteria in the motif enrichment analysis and we found the CTCF motifs to be highly enriched even with such a short input sequence (Figure 3b). The result demonstrates that Ntmax indeed denotes the first nucleotide of CTCF motif in a binding site.

### 3.4 Increasing the resolution of ChIP-seq data using DeepSNR

Chromatin immunoprecipitation (ChIP) has emerged as the most widely used assay over the last decade for genome-scale mapping of TF-DNA footprints and gene regulation. Although ChIP-seq method is an effective approach to decode regulatory relationships, it cannot resolve TF-DNA binding interactions at basepair resolution. Generally, size of the binding regions determined from ChIP-seq data are on the order of hundreds of base pairs. Taking advantage of single nucleotide resolution prediction capability of DeepSNR, the specificity of ChIP-seq data can be improved to base pair resolution. Notably, the deep learning model trained for one cell line of a particular TF is feasible to be applied on any other cell lines.

To exhibit the performance of DeepSNR on ChIP-seq data, we collected human CTCF ChIP-seq peak calling result for CD4 cell line published in (Martin et al., 2011). The dataset reports 20,272 peak regions each of which is 400 bp wide. When we applied DeepBind and DeepSNR concurrently on ChIP-seq data (see supplementary section S3), DeepBind confirmed CTCF binding in 11,750 peak regions (58% of all peaks). Hence, we restricted our further analysis to these peaks only. For each predicted binding site location by DeepSNR, we calculated the distance between Ntmax (described in previous section) and center of ChIP-seq binding region, the histogram of which is plotted in supplementary Fig S2. As evident, peak of the histogram is centered around −10 bp which implies that the first nucleotide of CTCF motif and therefore, the binding motif coincides with the ChIP-seq summit for most of the sites. Considering that CTCF motif is ∼20 bp long, center nucleotide of the motif detected by DeepSNR overlaps with ChIP-seq peak. However, for a significant number of ChIP-seq binding region, the binding motif (or binding site) is located far apart from ChIP-seq summit. This precise display of motif distribution within ChIP-seq peak region came into picture owing to the base pair resolution prediction of DeepSNR, which is otherwise not possible to visualize. Since, ChIP-seq summit doesn’t necessarily represent TF-DNA binding location as shown in Fig S2, DeepSNR can significantly reduce false positives/negatives in the analysis of ChIP-Seq data that results from consistently choosing the peak center as the putative TFBS. Besides, application of DeepSNR on ChIP-seq data delivers an unprecedented knowledge of the span of binding site which is not possible to attain using any motif search based methods.

**Table 1.**
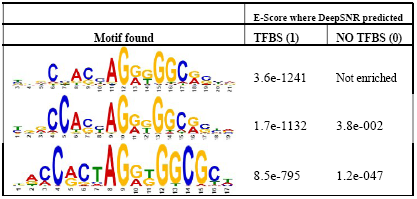
Motif enrichment analysis result of DeepSNR prediction on ChIP-seq data. CTCF motifs are significantly enriched where DeepSNR predicted binding site within ChIP-seq peak in comparison to locations where it didn’t.

To verify the credibility of DeepSNR prediction results on ChIP-Seq peaks, we performed independent motif enrichment analysis using nucleotide sequences where DeepSNR predicted ‘1’ and the sequences where it predicted ‘0’ within 400 bp wide ChIP-seq binding region. Table 1 shows the motifs derived from this analysis for CTCF. The striking difference in E-value (estimated statistical significance of a motif) between the CTCF motifs for positions where DeepSNR predicted protein-DNA binding (column 2) and those where it didn’t predict any binding (column 3) advocates for the efficacy of DeepSNR in pinpointing the binding location. While the CTCF motifs were identified as significantly enriched (E-values: 10-1132, 10-795) in DeepSNR predicted binding sequences, the enrichment scores were comparatively negligible in rest of the regions (E-values: 10-47, 10-2) demonstrating that DeepSNR is truly effective in predicting putative TF-DNA binding position. Thus,DeepBind and DeepSNR can be applied subsequently on ChIP-seq peak calling result to accomplish base pair resolution detection of TFBS which eventually leads to fewer false positive/negative rates.

**Fig. 4.**
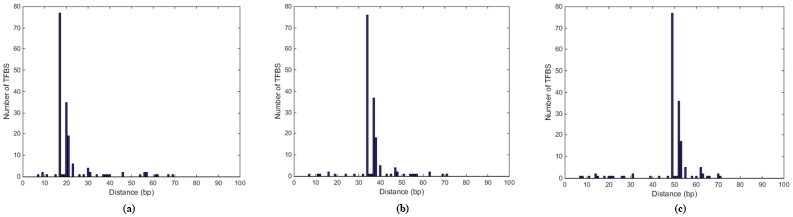
Histograms of distances of 168 experimentally verified 16 bp long regulatory sequences from Nt_max_ when they were concealed between (a) 27-42 bp, (b) 42-57 bp and (c) 58-73 bp. As evident, histogram peak follows the regulatory sequence validating that DeepSNR is very effective in discerning it from noisy background sequence.

### 3.5 DeepSNR recognizes functionally active regulatory sequence

Transcription factor (TF) proteins and DNA interacts with each other to regulate the transcription. One of the major impediments in unravelling the function of TF binding sites is to complement TFBS predictions with a high-throughput experimental approach that directly validates the functional contribution made by transcriptional regulatory motifs (Elnitski, Jin, Farnham, & Jones, 2006). In (Whitfield et al., 2012), the authors carried out a large-scale systematic functional analysis, at base-pair resolution, of predicted TF binding sites in four immortalized human cell lines (K562, HT1080, HCT116 and HepG2) by performing transient transfection assays on promoters.

There are 168 functionally verified 16 bp short regulatory sequences reported along with their genomic coordinates across four cell lines for CTCF. We wanted to investigate whether DeepSNR is sensitive enough to recognize these short controlling sequences when it is concealed inside 100 bp long genomic sequence. Therefore, combining adjacent nucleotides from human genome we extended each of these tiny sequences to the length of 100 bp for three instances such that they were placed in three different locations (27 - 42 bp, 42 - 57 bp and 58 – 73 bp). Next, we applied trained DeepSNR model of CTCF transcription factor on 168 sequences of each separate cases and recorded the nucleotide position of maximum score at the output of second max-pooling layer (Ntmax of previous section). The histogram of Ntmax across 168 sequences of each scenario are plotted in Fig. 4 depicting that DeepSNR responds actively to the change of locations of the most critical segment of input sequence required for TF regulation. The peak of histogram plots follows the positioning of the regulatory sequence which illustrates that DeepSNR is very accurate in recognizing the controlling sequence from noisy background sequence.

More intriguingly, the distance between histogram peak and the first nucleotide of regulatory sequence is 10 bp for all the cases which is reminiscent of the result discussed in previous section. In the process of determining transcriptional activity of a regulatory sequence the nucleotides making the greatest contribution to the TF-DNA binding were mutated such that it abolishes the binding (Whitfield et al., 2012) and transient transfection promoter activity assays were performed later on both wildtype and mutant sequences in order to determine substantial differences in transcriptional mechanism. It implies that the nucleotides pivotal for TF-DNA binding also govern the regulation orchestration of the transcription factor binding site. These nucleotides are eventually part of the binding motif and for CTCF, they are mostly located at 10 bp onwards within the motif sequence. Since, DeepSNR responds very sensitively to the positioning of regulatory (or motif) sequence, the model can play a role in the analysis of TF regulation scheme by locating the regulatory sequence in promoter region.

## 4 Conclusion

Computational prediction of transcription factor binding location from a genomic sequence remains a substantial challenge for the research community. While in previous decades genetic analyses focused on experimentally discovering TF-DNA binding (ChIP-seq, ChIP-exo etc.), due to the availability of deep sequencing the search using computational methods has meanwhile become a research focus. We developed DeepSNR, a deep learning framework to identify TFBS which performs better than sequence based approaches because it automatically learns the dependencies between nucleotides at different positions within the binding site description. DeepSNR is accomplished by successfully combining several technologies such as deconvNet, DeepBind and ChIP-exo that have been proved to achieve ground-breaking performance in their respective domain. The proposed model determines TFBS at base-pair resolution with high precision and recall which makes it suitable to discover regulatory sequences and to improve the specificity of ChIP-seq data. Currently, the state-of-the art methods for determining functional importance of TF utilize position weight matrices (PWMs) to identify regulatory (or motif) sequence as one of the initial procedures (Whitfield et al., 2012). It is not surprising that regulatory sequences derived using such technique exhibit highly different success rates in modelling TF-DNA binding. We have shown that DeepSNR responds very sensitively to the position of regulatory sequences when hidden at various places inside noisy background sequence. Therefore, instead of relying on PWM, DeepSNR can be applied to identify the location of regulatory sequence. In addition, this is the first application of deconvNet to address a computational biology problem.

Because of the limited availability of ChIP-exo data and to ensure wide applicability of DeepSNR, we focused on training the model with lone transcription factor on every occasion. However, predicting the binding location of multiple transcription factors simultaneously is an area that worth exploration in future. It can be expected that prediction of multiple TFs’ altogether might lead us to fully understand gene regulation and concurrent expression of genes as observed in expression array analysis.

## Supplementary Document for DeepSNR

### S1. Performance Evaluation Scheme

IoU is very popular in the field of pixel-level image segmentation since it discerns a proposed solution with respect to the ground truth in perceptually meaningful way. Precision (P) quantifies the correctness of a system i.e., the capacity to predict known binding sites accurately (true positives) while decreasing the number of wrongly predicted binding sites (false positives). Recall (R) measures the entirety, i.e., the ability of an algorithm to detect binding sites properly while minimizing the number of known binding sites that are erroneously missed (false negatives). Finally, F-Score (F), the harmonic mean of precision and recall, were calculated to have a measure of both methods’ efficiency in discovering putative TFBS. Mathematical definitions of these metrics are as follows:

IoU = TP / (TP + FN + FP)

Precision (P) = TP / (TP + FP)

Recall (R) = TP / (TP + FN)

F-Score = 2*P*R / (P + R)

### S2. Discovering the Most Significant Nucleotide

DeepBind’s mutation map scheme was deployed to evaluate the significance of each nucleotide on TF binding. Briefly, we mutated each nucleotide in a sequence with every possible point mutation and measured the change in affinity score for each mutation with respect to the affinity score of unaltered sequence using DeepBind. The average change in binding scores were evaluated for each base-pair and the nucleotide having the maximum average change in binding affinity was deemed to be the most important for TF binding.

### S3. Applying DeepBind and DeepSNR concurrently on ChIP-seq data

The DeepSNR model for CTCF transcription factor was trained using 100 bp long DNA sequences from ChIP-exo data of HeLa cell line. Therefore, to apply trained DeepSNR model on 400 bp wide ChIP-seq binding region, we divided the entire region into 5 bins of 100 bp each using a sliding window of 75 bp. Next, we fed each bin into DeepBind to determine the presence of putative binding site in the input sequence. If DeepBind confirms TF-DNA binding in a bin then we applied DeepSNR to predict the binding location in that bin. In case of DeepBind predicting no binding, we moved to the next bin to repeat the same process and finally the binary outcome through DeepSNR for all the bins were combined as OR operation to achieve the base pair resolution prediction result of whole ChIP-seq binding region. DeepSNR was employed in conjunction with DeepBind because DeepSNR is trained such that its input sequence is known to contain a binding site. Therefore, applying DeepSNR on a sequence that doesn’t contain binding site may lead to erroneous result.

### Supplementary Tables and Figures

**Table S1.** Information of dataset used for predicting binding location of Androgen Receptor (AR) and Glucocorticoid Receptor (GR)

| TF name | Total no of samples<br>(training,validation,testing) | Binding<br>length | Input sequence<br>length | Source |
| --- | --- | --- | --- | --- |
| Androgen receptor (AR) | 402472 (272472, 30000, 100000) | 58 bp | 100 bp | GSE43785 |
| Glucocorticoid receptor (GR) | 28865 (22865, 1000, 5000) | 79 bp | 200 bp | GSE79432 |

**Table S2.** Performances of DeepSNR and MatInspector in predicting AR and GR binding sites

| TF name | Method | Median precision | Median recall | Median F1-score | Median IoU |
| --- | --- | --- | --- | --- | --- |
| AR | DeepSNR | 0.871 | 0.931 | 0.9 | 0.812 |
|  | MatInspector | 0.894 | 0.327 | 0.493 | 0.327 |
| GR | DeepSNR | 0.507 | 1.0 | 0.669 | 0.503 |
|  | MatInspector | 0.482 | 0.177 | 0.152 | 0.265 |

**Figure S1.**
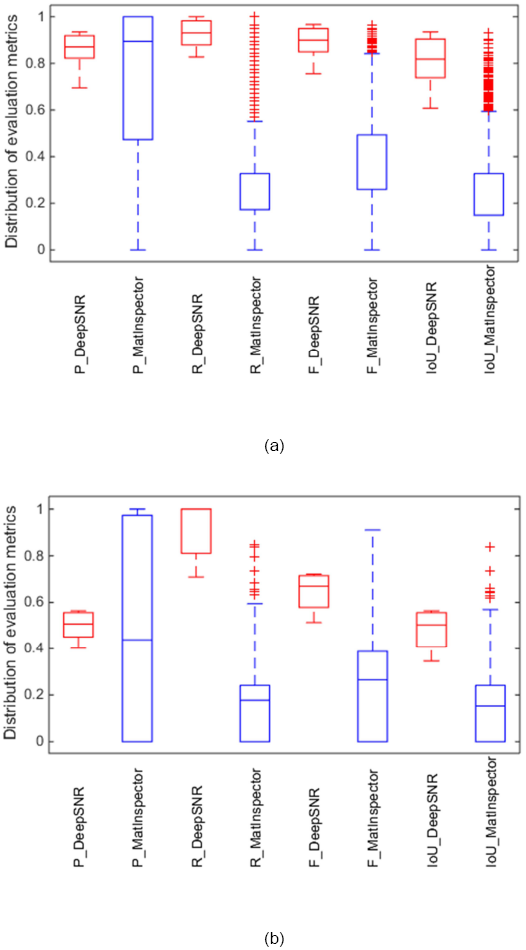
Similar to CTCF transcription factor, DeepSNR has shown better performance than MatInspector in predicting binding locations of (a) AR and (b) GR transcription factors.

**Figure S2.**
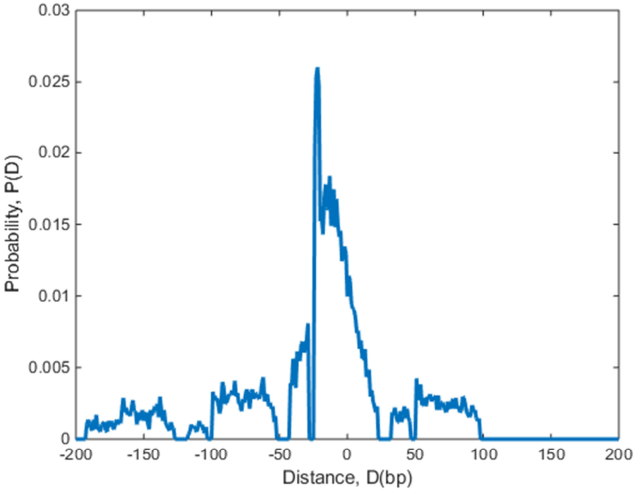
Probability distribution of distance between CTCF motif (determined by DeepSNR) and ChIP-seq peak centre

